# Synthetic Antigen-Presenting Cells for Adoptive T Cell Therapy

**DOI:** 10.1101/2021.02.01.429167

**Authors:** Shreyas N. Dahotre, Anna M. Romanov, Fang-Yi Su, Gabriel A. Kwong

## Abstract

Adoptive T cell therapies are transforming the treatment of solid and liquid tumors, yet their widespread adoption is limited in part by the challenge of generating functional cells. T cell activation and expansion using conventional antigen-presenting cells (APCs) is unreliable due to the variable quality of donor-derived APCs. As a result, engineered approaches using nanomaterials presenting T cell activation signals are a promising alternative due to their ability to be robustly manufactured with precise control over stimulation cues. In this work, we design synthetic APCs that consist of liposomes surface-functionalized with peptide-major histocompatibility complexes (pMHC). Synthetic APCs selectively target and activate antigen-specific T cell populations to levels similar to conventional protocols using non-specific αCD3 and αCD28 antibodies without the need for costimulation signals. T cells treated with synthetic APCs produce effector cytokines and demonstrate cytotoxic activity when co-cultured with tumor cells presenting target antigen *in vitro*. Following adoptive transfer into tumor-bearing mice, activated cells control tumor growth and improve overall survival compared to untreated mice. Synthetic APCs could potentially be used in the future to improve the accessibility of adoptive T cell therapies by removing the need for conventional APCs during manufacturing.

## 1. Introduction

Adoptive cell transfer (ACT) of autologous tumor-infiltrating lymphocytes (TILs) is a cell-based therapy that has garnered increasing interest for treating cancer. For example, in patients with metastatic melanoma, overall response rates ranging from 40-70% have been observed, with as much as 20% of cases resulting in complete or durable regression.^[1–2]^ Despite its therapeutic potential, ACT has yet to become the standard-of-care for cancer treatment in part due to the complexity of generating sufficient numbers of highly functional TILs. This multi-step pipeline begins with isolation of TILs from patient tumor biopsies, an initial TIL outgrowth phase of ~2 weeks, and a rapid expansion phase during which TILs are stimulated using autologous or allogenic antigen-presenting cells (APCs) to reach large numbers (10^10^–10^11^ cells).^[3]^ The expanded TILs are transferred into patients and supported by co-infusions of high-dose interleukin-2 (IL-2). One limitation of this process is the need for patient APCs, which are primarily mature dendritic cells (DCs) derived from patient peripheral blood mononuclear cells (PBMCs). This process requires complex culture protocols and can still result in inadequate TIL activation, as DCs from cancer patients often have an impaired ability to present tumor antigens and properly activate antigen-specific T cells.^[4–5]^

Engineered alternatives to natural APCs for *ex vivo* TIL activation include both cell-based and biomaterials-based approaches. K562 cells engineered to display αCD3 and αCD28 antibodies or scFvs on their surface have previously been shown to activate and expand T cells by greater than 50-fold over the course of 9 days.^[6–8]^ For clinical use, GMP-grade magnetic beads (e.g., Dynabeads) or polymeric matrices (e.g., Miltenyi T Cell TransAct^™^) coupled with αCD3 and αCD28 antibodies are routinely used to activate T cells prior to ACT.^[9–10]^ These approaches are adequate for therapies involving genetically engineered transgenic T cells with T cell receptors (TCR) or chimeric antigen receptors (CAR) in which the cell product consists of a single antigen-specific population. However, for TIL therapy, nonspecific stimulation by αCD3/αCD28 results in the activation of both tumor-specific and non-specific clones, leading to an overall decrease in antigen-specificity and limiting the ability of tumor-specific T cells to persist post-transfusion.^[11–13]^ More sophisticated strategies for selective activation and expansion of antigen-specific T cells involve the use of magnetic nanoparticles^[13–16]^ or 3D-scaffolds^[17–18]^ decorated with major histocompatibility complexes (MHCs) presenting disease-specific peptides. These pMHC-based materials have been shown to be able to rapidly activate and expand low-frequency antigen-specific T cell populations directly from mouse splenocytes or human PBMCs, producing cells capable of potent cytotoxic activity in pre-clinical models.^[13-14, 17-18]^

Recombinant pMHC molecules are typically used as multimeric complexes to analyze or sort antigen-specific T cells by flow-, mass-, or DNA-based cytometry.^[19–23]^ Here we develop synthetic antigen-presenting cells (synAPCs) comprising liposomes surface-conjugated with pMHC molecules for antigen-specific activation of T cells. We optimize pMHC density on liposomes for T cell binding and demonstrate that synAPCs activate antigen-specific T cells *in vitro*. We show that T cells activated via synAPCs produce effector cytokines and are capable of killing target cells when co-cultured with tumor cells presenting a cognate antigen. When adoptively transferred to mice bearing B16-OVA tumors, OVA-specific T cells stimulated by synAPCs control tumor growth and extend overall survival. Our results support the development of synthetic APCs for use in adoptive immunotherapy with antigen-specific T cells.

## 2. Results and Discussion

### 2.1. Binding of Synthetic APCs to T Cells Depends on pMHC Surface Density

Previous work using pMHC multimers and pMHC-coupled nanoparticles demonstrated that pMHC ligand density impacts T cell binding and activation.^[15, 24-25]^ We therefore first optimized the surface display density of pMHC molecules on synthetic APCs to maximize T cell binding. We formulated liposomes using a fixed 6:4 molar ratio of DSPC:cholesterol as the basis for synAPCs as the fluid bilayer enables reorganization of pMHC into microclusters to improve T cell stimulation during immune synapse formation^[26–27]^. We further varied the molar percentage of DSPE-PEG-maleimide in the liposomes from 0% to 1% to provide a bioconjugation strategy to attach and titrate pMHC molecules onto the surface (**Figure 1A, B**). To couple pMHC to the exposed maleimide groups, we reacted liposomes with pMHC refolded using heavy chains engineered with a C-terminal cysteine to introduce a free thiol group that could be selectively reduced for bioconjugation. The use of an engineered terminal cysteine residue minimizes disruption of native disulfide bonds^[28]^ and retains proper orientation of the protein for recognition by TCRs.^[29]^ We found that all lipid compositions formed particles with an average diameter less than 200 nm and a polydispersity index (PDI) ranging from 0.08 to 0.21 as measured by dynamic light scattering (DLS) (**Figure 1B**). After reaction with thiolated pMHC, the size of the liposomes increased (136 nm to 268 nm, 127 nm to 139 nm, and 143 nm to 183 nm) for all molar percentages tested (0.1%, 0.5%, and 1% DSPE-PEG-maleimide, respectively) (**Figure 1B**). We then measured pMHC concentration on reacted liposomes using UV-vis spectroscopy and found that the pMHC coupling efficiency (i.e., the final pMHC:DSPE-PEG-maleimide ratio) increased from 0.13 to 0.33 as DSPE-PEG-maleimide increased in our formulations (**Figure 1B**). These pMHC-labelled liposomes were stable with minimal size change observed after storage for 1 month at 4°C (**Figure 1C**). To identify the formulation that maximized binding to T cells, we reacted liposomes containing 1% molar ratio of the lipophilic dye DiO and 0, 0.1, 0.5, or 1.0% molar ratios of DSPE-PEG-maleimide with Db-GP33 pMHC which presents a peptide epitope derived from lymphocytic choriomeningitis virus (LCMV). We then incubated functionalized liposomes with splenocytes from P14 transgenic mice, whose CD8+ T cells express a TCR specific for the LCMV GP33 antigen, and measured DiO signal on T cells by flow cytometry across a range of synAPC concentrations (0-40 μg/ml). Fitting the data to a one-site model of receptor-ligand binding, we compared asymptotic maximum DiO signal for 0.1, 0.5, and 1.0% DSPE-PEG-maleimide liposomes with control 0% liposomes and found that a 0.5% DSPE-PEG-maleimide molar percentage resulted in the greatest change in MFI (93-fold, 289-fold, and 223-fold increase for 0.1, 0.5, and 1.0%, respectively) (**Figure 1D**). This formulation of synAPCs was used for subsequent experiments.

**Figure 1.**
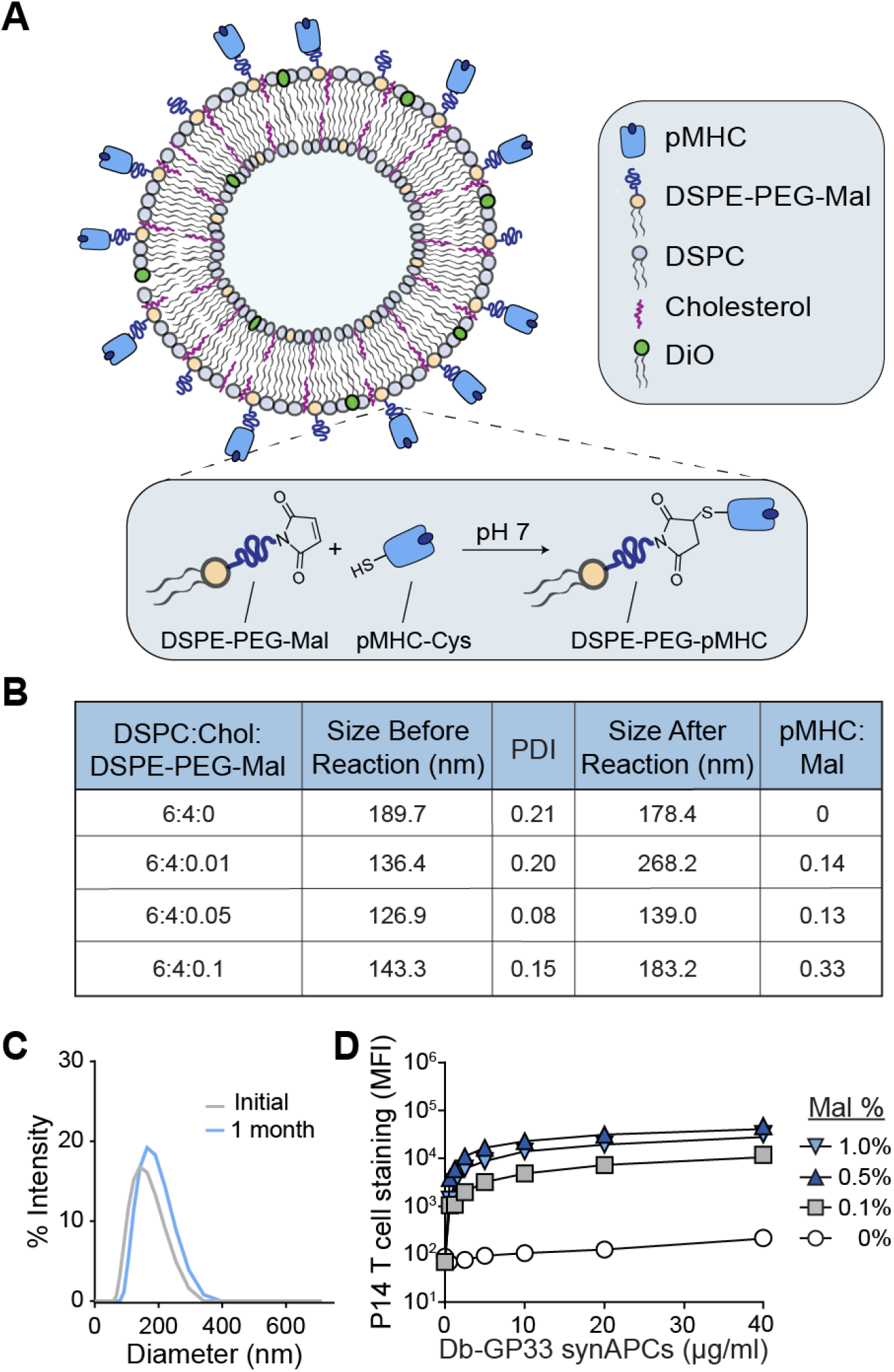
The surface density of pMHC on synthetic APCs impacts T cell binding. (A) SynAPCs are formulated with DSPC, cholesterol, and DSPE-PEG-maleimide. pMHC engineered with a C-terminal cysteine residue reacts with free maleimide on the liposome surface. (B) Table summarizing liposome characteristics for various DSPE-PEG-maleimide ratios. (C) DLS of synAPCs showing that liposomes remain stable for at least 1 month at 4°C. (D) Binding curve of pMHC targeted synAPCs staining target T cells as quantified as MFI by flow cytometry. A 0.5% DSPE-PEG-maleimide ratio was chosen for subsequent experiments. Data fit to a one-site model of receptor-ligand binding.

### 2.2. Synthetic APCs Bind to T Cells in an Antigen-Specific Manner

We next sought to demonstrate that synthetic APCs selectively bind T cells in an antigen-specific manner. We coupled Db-GP33 pMHC to liposomes containing 1% DiO and incubated them with splenocytes from either P14 mice or pmel control mice – a strain whose CD8+ T cells recognize an irrelevant human gp100 antigen (**Figure 2A**). We found that Db-GP33 synAPCs bound to nearly all (88.1%) P14 CD8+ T cells with minimal background staining (3.45%) to pmel CD8+ T cells (**Figure 2B**). To confirm that this binding was mediated by pMHC and not through non-specific interactions with the liposome surface, we incubated P14 and pmel CD8+ T cells with bare liposomes lacking pMHC. We found that there was no significant difference in binding to pmel T cells between Db-GP33 synAPCs and bare liposomes while binding to P14 T cells was significantly increased using synAPCs conjugated with Db-GP33 pMHC (****p<0.001 by one-way ANOVA) (**Figure 2C**). In a competition assay using fluorescent Db-GP33 synAPCs incubated with a mixture of P14 and pmel CD8+ T cells, we found that Db-GP33 synthetic APCs preferentially bound to P14 T cells compared to pmel T cells (95.2% vs. 4.8% liposome positive, respectively) (**Figure 2D, E**). Together, these results demonstrate that pMHC labeling is necessary for synAPCs to target and discriminate between antigen-specific CD8+ T cell populations.

**Figure 2.**
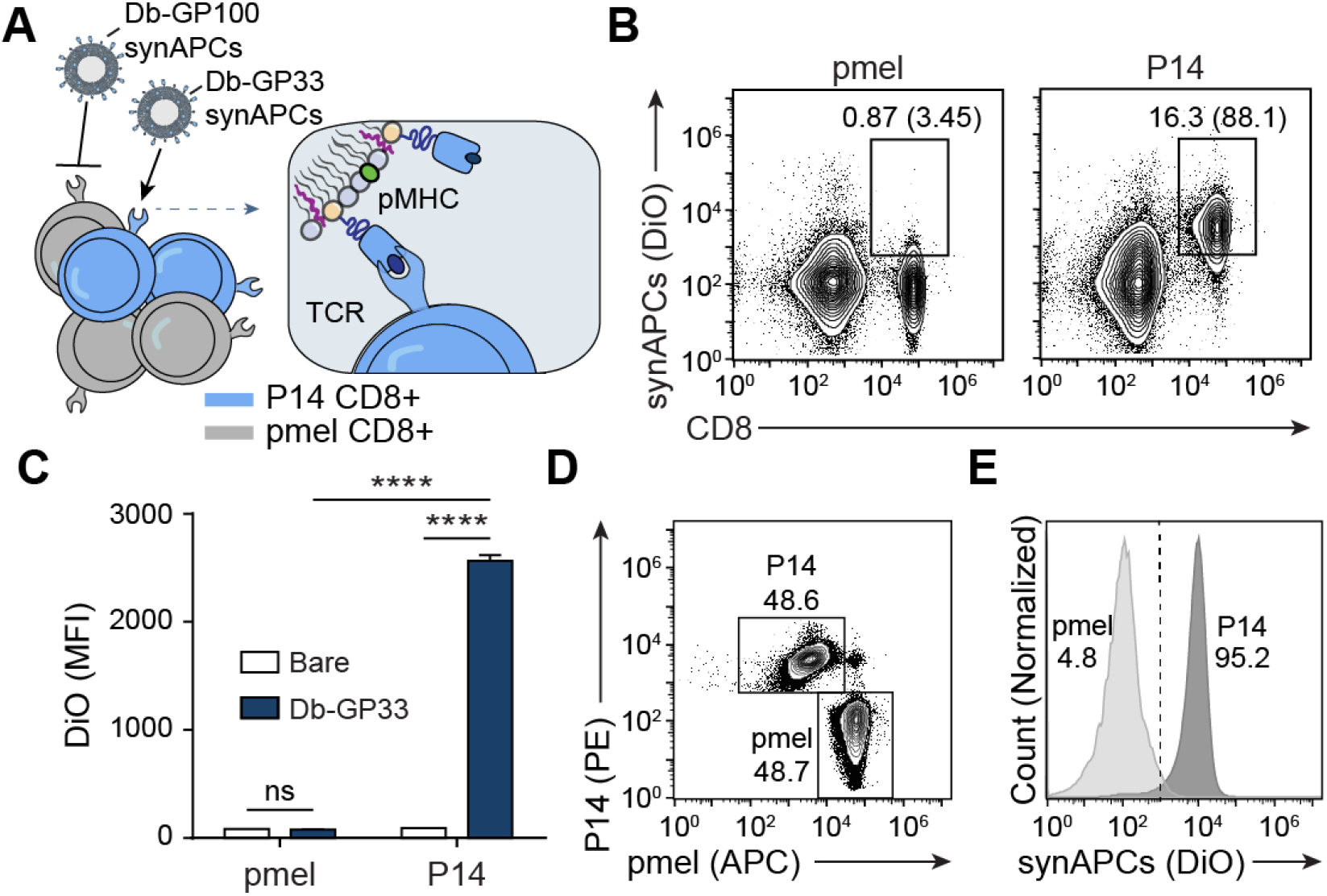
Synthetic APCs bind to T cells in an antigen-specific manner. (A) Db-GP33 synAPCs are incubated with CD8+ T cells from P14 and control pmel TCR transgenic mice. (B) Flow plots of synAPC binding to CD8+ T cells from pmel or P14 TCR transgenic mice. Frequencies depicted are based on gating on total splenocytes and gating on CD8+ cells. (C) MFI of CD8+ cells from pmel or P14 transgenic mice stained with untargeted or Db-GP33 synAPCs. Data shown as mean ± s.d. n=3 ****p<0.0001 by two-way ANOVA with Tukey’s multiple comparison test. (D) Flow plot of pmel and P14 CD8+ T cell mixture used in competition assay. pmel and P14 cells were pre-stained with αCD8-APC and αCD8-PE, respectively, prior to combining. (E) Flow plot of competition assay in which a mixture of pmel and P14 CD8+ T cells are incubated with Db-GP33 synAPCs.

### 2.3. CD8+ T Cells are Activated by Synthetic APCs Without Costimulation

We next asked if synthetic APCs activate naïve CD8+ T cells, as soluble pMHC tetramers have previously been shown to activate CD8+ T cells even in the absence of costimulatory signals (**Figure 3A**).^[24, 30]^ We therefore compared activation of P14 CD8+ T cells by synAPCs presenting Db-GP33 against activation by αCD3/αCD28 antibodies for 24 hours. αCD3 and αCD28 antibodies activate T cells by TCR stimulation and costimulatory signaling respectively and are the standard for *ex vivo* activation of T cells prior to viral transduction and adoptive cell transfer in CAR T cell therapies.^[31]^ Compared to naïve cells, we found that P14 CD8+ T cells incubated with Db-GP33 synAPCs showed increased expression of the activation markers CD44 and CD69 to levels that were equivalent to cells treated with αCD3/αCD28 antibodies (**Figure 3B**). Upregulation of activation markers was seen even in the absence of costimulatory signaling with synAPCs, demonstrating that potent stimulation of the TCR through multivalent presentation of pMHC molecules on the surface of a lipid membrane surface is sufficient to induce T cell activation. To verify that activated cells exhibited cytotoxic activity, we treated P14 CD8+ T cells with bare liposomes (lacking pMHC), Db-GP33 synAPCs, or αCD3/αCD28 and then co-cultured these T cells with MC38 tumor cells alone or MC38 tumor cells pulsed with GP33 peptide. We observed no significant difference in the expression of IFNγ by intracellular staining when T cells treated with bare liposomes were co-cultured with MC38 WT or MC38-GP33 cells (**Figure 3C**). By contrast, T cells that were stimulated with Db-GP33 synAPCs showed a significant increase in IFNγ expression when cultured with GP33-pulsed MC38 cells (****p<0.001 by two-way ANOVA). To verify that activation by synAPCs leads to target cell killing, we used a second TCR/pMHC combination and primed CD8+ T cells from OT1 TCR transgenic mice, which recognize a Kb-restricted ovalbumin antigen (OVA), with bare liposomes,, Db-GP33 synAPCs, or Kb-OVA synAPCs before co-culturing each population with B16-OVA tumor cells. We found that only OT1 T cells that had been stimulated with Kb-OVA synAPCs killed B16-OVA cells with increasing effector:target cell ratios (**Figure 3D**). These results show that pMHC-conjugated liposomes activate antigen-specific T cell populations *in vitro* and that activated cells retain cytotoxic activity.

**Figure 3.**
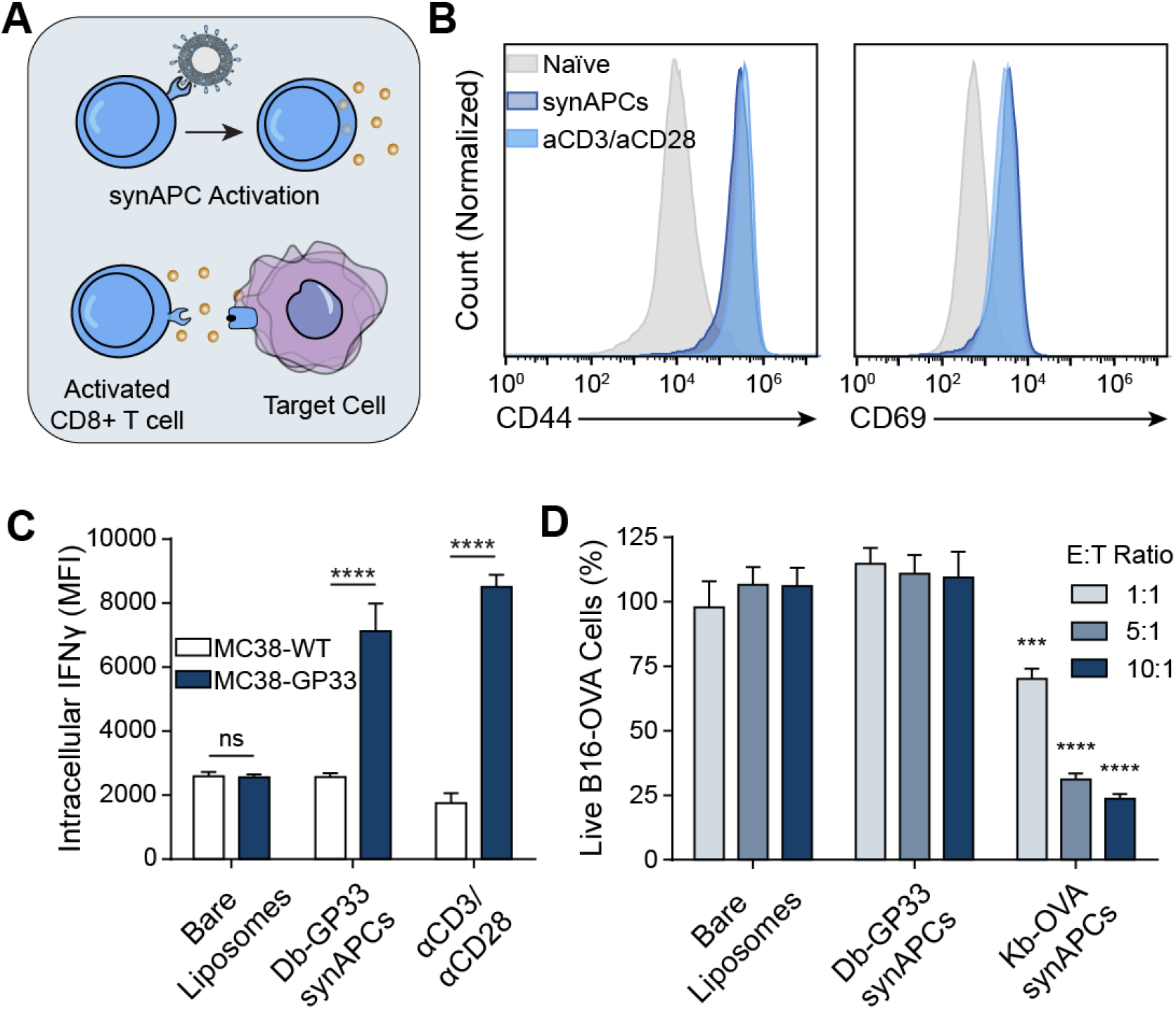
T cells activated by synAPCs *in vitro* demonstrate cytotoxic activity. (A) pMHC liposomes stimulate T cells through pMHC-TCR interactions. Activated T cells then exert cytotoxic effects on target cells. (B) Histograms of CD44 and CD69 expression on T cells stimulated with synAPCs or anti-CD3/anti-CD28 antibodies. (C) P14 CD8+ T cells activated with Db-GP33 synAPCs have higher IFNγ expression when incubated with target cells pulsed with GP33 peptide than when incubated with control cells. Data shown as mean ± s.d. n=3 ****p<0.0001 by two-way ANOVA with Tukey’s multiple comparison test. (D) OT1 CD8+ T cells stimulated with Kb-OVA synAPCs show cytotoxic activity against B16-OVA target cells. Data shown as mean ± s.d. n=4 ***p<0.001, ****p<0.0001 by two-way ANOVA with Tukey’s multiple comparison test. Significance relative to untargeted controls at same E:T ratio.

### 2.4. Synthetic APCs are Internalized by Antigen-Specific T Cells at Physiological Temperatures

Previous work demonstrated that pMHC multimers are rapidly internalized by T cells through TCR clustering and receptor-mediated endocytosis after incubation at physiological temperatures.^[32]^ We therefore probed the fate of synthetic APCs after engaging cognate T cells at 37°C. We synthesized Db-GP33 synAPCs labeled with DiO, incubated them with P14 and control pmel CD8+ T cells at 4 or 37°C, and imaged cells by confocal microscopy. We observed DiO signals largely localized to the periphery of P14 T cells incubated at 4°C. By contrast, cells stained at 37°C had punctate spots of DiO signal, indicating synAPC internalization and clustering (**Figure 4A**). To corroborate this finding, we analyzed T cells stained with DiO-labeled synAPCs at 4 or 37°C by flow cytometry. We then treated stained cells with an acidic wash buffer to strip cell surface proteins and reanalyzed DiO fluorescence by flow cytometry. We found that DiO fluorescence decreased for cells incubated at 4°C, indicating that synAPCs remained on the cell surface (**Figure 4B**). For cells treated at 37°C, however, we observed no change in fluorescence, demonstrating that synAPCs are efficiently internalized by T cells (**Figure 4B**). To track the migration of internalized synAPCs through the endocytic pathway, we stained T cells with LysoTracker Red dye to label lysosomes and incubated stained cells with DiO-labeled synAPCs. We fixed cells 0.5, 2, and 4 hours after synAPC addition and imaged cells by confocal microscopy. We observed punctate spots inside cells within 30 minutes, demonstrating that synAPCs are rapidly internalized, and found that synAPCs co-localized with lysosomes by 4 hours (**Figure 4C**). These results demonstrate that synAPCs are taken up by T cells at physiological temperatures.

**Figure 4.**
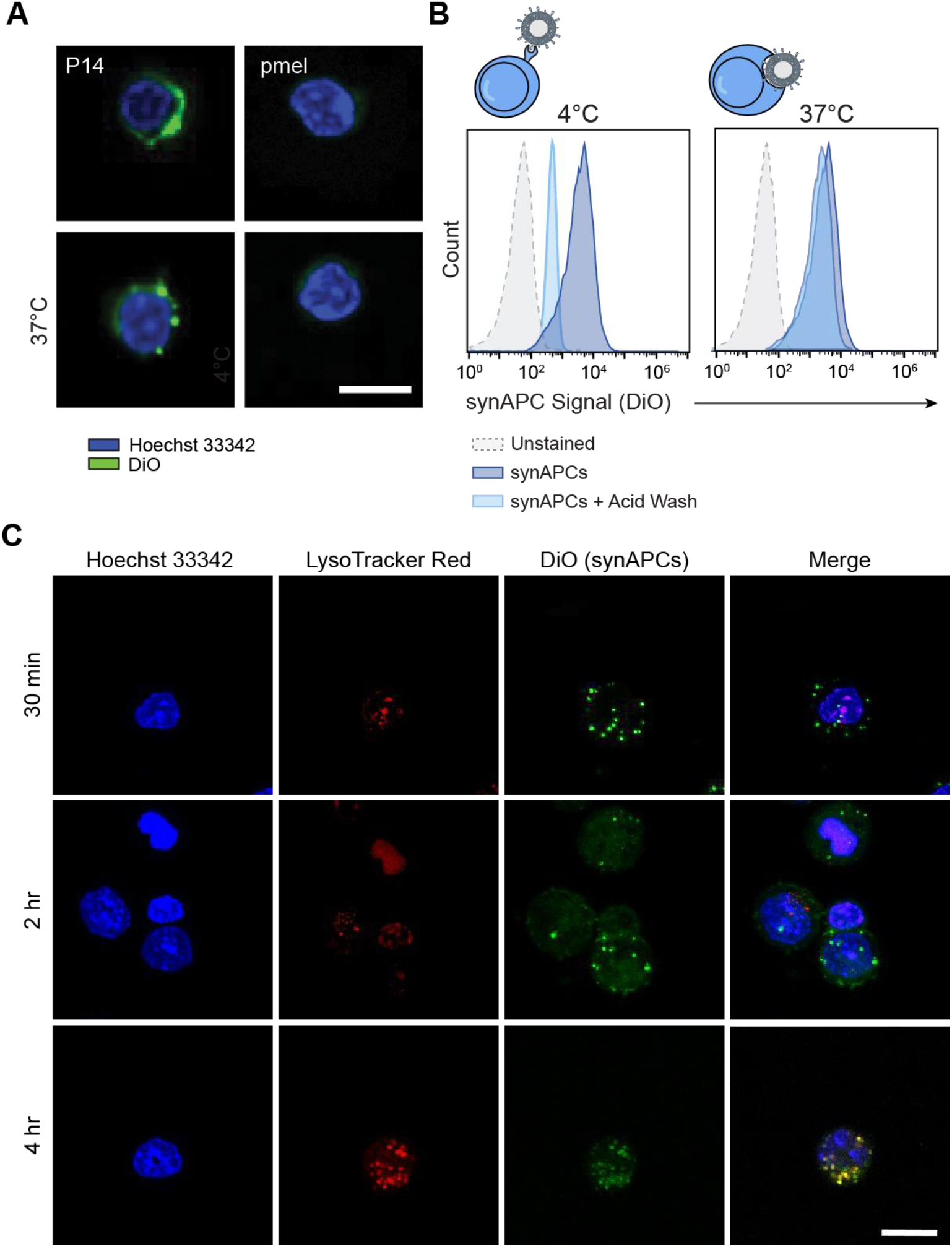
Synthetic APCs are rapidly internalized by antigen-specific T cells. (A) Representative images of P14 and pmel CD8+ T cells stained with DiO-labeled Db-GP33 synAPCs at 4 or 37°C. Scale bar = 15 μm. (B) Representative histograms for cells stained with synAPCs at 4 or 37°C and analyzed by flow cytometry before and after treatment with an acid wash to strip cell surface receptors. (C) Representative images of CD8+ T cells labeled with a lysosome dye and stained with synAPCs for various lengths of time. Cells were fixed at the times indicated and imaged by confocal microscopy. Scale bar = 15 μm.

### 2.5. Antigen-Specific T Cells Activated by Synthetic APCs Control Tumor Growth After ACT

To determine if antigen-specific T cells activated by synAPCs could be used for adoptive immunotherapy, we stimulated OT1 CD8+ T cells with Kb-OVA synAPCs for 24 hours prior to adoptive transfer into mice bearing well-established, syngeneic B16-OVA tumors (mean tumor size at ACT was ~40 mm^3^, n=7 per group) (**Figure 5A**). We transferred 2×10^5^ activated cells per mouse and supplemented ACT with multiple doses of IL-2 as TIL therapies in the clinic include infusions of high dose IL-2 to improve the proliferation of adoptively transferred cells.^[33–34]^ Compared to untreated mice, we found that mice receiving T cell therapy controlled tumor outgrowth, resulting in significantly smaller tumors (*p<0.05 bv two-way ANOVA) (**Figure 5B**) and a significantly increased survival time (**Figure 5C**). These results demonstrate that pMHC liposomes generate functional cells capable of exerting cytotoxic effects *in vivo*.

**Figure 5.**
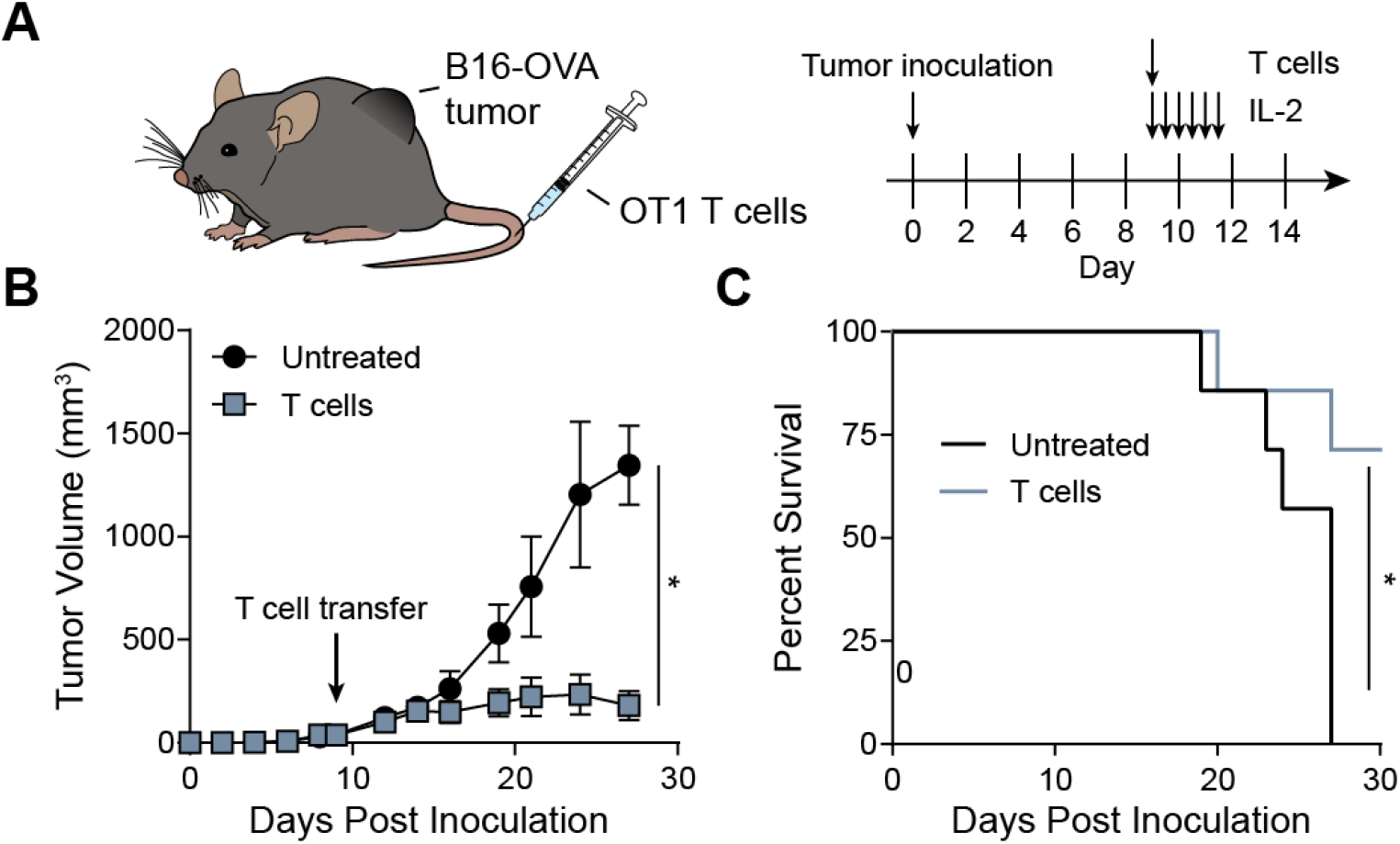
Adoptive transfer of T cells activated by synthetic APCs controls tumor growth. (A) Timeline of ACT experiment. (B) Tumor growth curves for mice with and without treatment. Data shown as mean ± s.d. n=7 *p<0.05 by two-way ANOVA. (C) Survival curve for mice without and without treatment. *p<0.05 by log-rank test.

## 3. Conclusion

Adoptive transfer of TILs is a promising therapy for cancer, but our ability to effectively activate and expand sufficient numbers of TILs for treatment is limited by the variable quality of autologous APCs. To improve the accessibility of ACT therapies, new approaches using synthetic materials that allow for precise control of activation signals are being developed. In this work, we design and formulate synthetic APCs consisting of liposomes decorated with pMHC molecules to activate antigen-specific T cells for adoptive cell therapy. We optimize pMHC ligand density to maximize binding to T cells and show that T cells specifically interact with synAPCs presenting target pMHC. *In vitro*, antigen-specific T cells cultured with synAPCs rapidly increase the expression of the activation markers CD69 and CD44, and OT1 T cells activated by this approach exhibit killing activity when co-cultured with B16-OVA tumor cells. When adoptively transferred into B16-OVA tumor bearing-mice, we find that OT1 T cells initially activated *ex vivo* with synAPCs effectively control tumor burden and improve survival time.

Here we show that pMHC stimulation alone using synAPCs is sufficient for T cell activation and therapy. However, future improvements of synAPCs may include the addition of costimulatory or cytokine signals such as IL-7/IL-15 which have previously been shown to skew expanded cells towards Tcm and Tscm phenotypes that improve the persistence of adoptively transferred cells *in vivo*.^[35]^ Additionally, our observation that synAPCs are readily internalized at physiological temperatures may allow for the modulation of antigen-specific T cells by delivery of small molecule drugs encapsulated in pMHC-functionalized liposomes. This would represent an advance in the precision of T cell delivery methods as previous approaches have primarily targeted broad T cell subsets.^[36–38]^ Last, although the identity of tumor neoantigens is difficult and patient specific, the development of large yeast-display libraries has improved our ability to identify the peptide targets of TILs and orphan T cells with unknown antigen specificity.^[39]^ In the future, we envision that libraries of synthetic APCs presenting neoepitopes can be generated by rapid pMHC exchange strategies^[40]^ to activate and expand neoantigen-specific T cells for personalized immunotherapy.

## 4. Experimental Methods

### 4.1. Liposome Synthesis and pMHC Conjugation

All lipids and extrusion supplies were purchased from Avanti Polar Lipids. A lipid solution consisting of 1,2-distearoyl-sn-glycero-3-phosphocholine (DSPC), cholesterol, and 1,2-distearoyl-sn-glycero-3-phosphoethanolamine-N-[maleimide(polyethylene glycol)-2000 (DSPE-PEG2000-maleimide) at a 59.5/40/0.5 molar ratio was dried under nitrogen and placed in vacuum chamber overnight to form a thin film. For liposome binding and internalization experiments, 1% DiO (Invitrogen) was added to the lipid mix prior to drying. Lipids were rehydrated in PBS at 2 mg/ml in a 60°C water bath for 30 mins, vortexed on high speed for 1 minute, and sonicated in an ultrasonic bath (Branson) for 15 minutes, or until no clumps were visible. After sonication, the liposomes were extruded ten times each through 0.8, 0.4, 0.2, and 0.1 μm polycarbonate membranes. Particle size and polydispersity were measured by dynamic light scattering (Malvern Zetasizer Nano ZS).

To generate pMHC molecules for bioconjugation, codon-optimized gBlocks for H2-Db, H2-Kb, and human β2m were purchased from IDT and cloned into pET3a vectors (Novagen). H2-Db and H2-Kb genes were engineered with a C-terminal cysteine by site-directed mutagenesis (NEB), and pMHC molecules were expressed and refolded as described previously^[40]^. Refolded Cys-terminated pMHC monomers were reduced with TCEP at a 3:1 TCEP:pMHC molar ratio for 2 hr at 37°C and then mixed with liposomes overnight at room temperature at a 2:1 pMHC:maleimide molar ratio. Unreacted pMHC was removed by buffer exchange into PBS using Amicon 100K spin filters. Final lipid concentration was measured using a Phospholipid Assay Kit (Sigma) with unreacted liposomes used to generate the standard curve. The concentration of conjugated pMHC was determined by measuring A280 using unreacted liposomes as a blank.

### 4.2. Primary T cell Isolation and Activation

Spleens from P14, pmel, or OT1 TCR transgenic mice (Jackson Labs) were dissociated in R10 media (RPMI 1640 (Gibco) + 10% FBS (Gibco) + 1% pen/strep (Gibco)), and red blood cells were lysed using RBC Lysis Buffer (Biolegend). CD8+ T cells were isolated using a CD8a+ T Cell Isolation Kit (Miltenyi Biotec). For pMHC liposome activation *in vitro*, T cells were cultured at 2×10^6^ cells/ml in T cell media (R10 supplemented with 1X non-essential amino acids (Gibco) + 1 mM sodium pyruvate (Gibco) + 0.05 mM 2-mercaptoethanol (Sigma)), and liposomes were added at 1 μg/ml. For antibody-activated controls, T cells were cultured in T cell media supplemented with soluble anti-mouse CD28 (2 μg/ml, Clone: 37.51, BD Pharmingen) and 30 U/ml rhIL-2 (Roche) at 2×10^6^ cells/ml in wells coated with anti-mouse CD3e (1 μg/ml, Clone: 145-2C11, BD Pharmingen).

### 4.3. In vitro Binding Affinity and Staining

Liposomes containing 1% DiO were formed with various ratios of DSPE-PEG2000-Maleimide (0, 0.1, 0.5, 1.0%) and reacted with Cys-terminated pMHC as described above. Naïve T cells (1×10^6^ cells/sample) were stained with 0-40 μg/ml pMHC liposomes in FACS buffer (1X DPBS + 2% FBS + 1mM EDTA + 25mM HEPES) for 30 min on ice. Cells were washed 4 times with 1 ml FACS buffer before analysis on a BD Accuri C6. Titration curves from median MFI were fit to one-site specific binding curve to determine the apparent dissociation constant (K^d,app^).

For validation of antigen-specific binding, 1×10^6^ P14 or pmel splenocytes were co-stained with Db-GP33 liposomes (10 μg/ml) and anti-mouse CD8-PE (Biolegend, Clone: 53-6.7) for 30 min on ice. Cells were washed 3 times and analyzed on a BD Accuri C6. For competition experiments, P14 and pmel CD8^+^ cells were pre-stained with anti-mouse CD8-PE and anti-mouse CD8-APC, respectively, washed thoroughly with FACS buffer, and mixed together (1:1) in the same tube. The cells were stained with Db-GP33 liposomes as described above and analyzed on a BD Accuri C6. Gates were drawn around PE+ and APC+ cells and were used to quantify liposome uptake (DiO) in each group.

### 4.4. T Cell Killing Assay with Pulsed MC38 Cells

P14 CD8+ T cells were isolated and activated by pMHC liposomes or anti-CD3/anti-CD28 as described above. The media was changed and supplemented with 30 U/ml rhIL-2 after 48 hr. On day 5 post-activation, MC38 target cells were pulsed with GP33 peptide in OptiPRO SFM media (Gibco) for 1 hr at 37°C. Pulsed MC38 cells were washed 5 times with PBS, seeded in a 96 well plate at 6×10^5^ cells/ml in T cell media, and incubated for 4 hr to allow adhesion. Activated P14 T cells were washed 5 times with PBS and incubated with MC38 cells at a 5:1 effector-to-target ratio overnight. For analysis of IFNγ expression, cells were washed with PBS, blocked with Fc Shield (Tonbo) for 10 minutes on ice, and stained with anti-mouse CD8 (Biolegend, Clone: 53-6.7) and anti-mouse IFNγ (Biolegend, Clone: XMG1.2) for 30 min on ice before collection on a BD Accuri C6.

### 4.5. T Cell Killing Assay with B16-OVA Cells

OT1 CD8+ T cells were isolated as described above and incubated with unconjugated, Db-GP33, or Kb-OVA liposomes at 1 μg/ml in T cell media for 24 hr. T cells were washed and then co-incubated with Renilla luciferase expressing B16-OVA cells (~50-60% confluency) at 1:1, 5:1, or 10:1 effector-to-target cell ratios in T cell media for 24 hr. Luciferase activity was measured by adding ViviRen^™^ In Vivo Renilla Luciferase Substrate (Promega) according to the manufacturer’s instructions and collecting on a Cytation 5 plate reader (BioTek).

### 4.6. Quantification of Liposome Internalization

P14 CD8+ T cells were isolated as described above and incubated with DiO-labeled Db-GP33 liposomes and anti-mouse CD8 (Biolegend, Clone: 53-6.7) at 4°C or 37°C for 30 min. Cells were washed with FACS buffer, and a portion of stained cells were analyzed on a BD Accuri C6. The remaining cells were incubated in an acid wash solution (0.5 M NaCl + 0.5 M acetic acid, pH 2.5) for 5 min to strip cell surface proteins as described previously^[41]^ before reanalysis on a BD Accuri C6.

### 4.7. Confocal Microscopy

Glass cover slips were coated with poly-L-lysine for 2 hours at 37°C prior to use. 2×10^6^ P14 CD8+ T cells were first stained with 250 nM Lysotracker Red DND-99 (Invitrogen) for 30 minutes, washed with PBS, and then stained with 1 μg/ml Db-GP33 liposomes (containing DiO) at 37°C. At various time points, cells were collected and washed 4 times with PBS, fixed with 4% PFA, and stained with Hoechst 33342 (Invitrogen). Samples were spun onto cover slips at 1000xg for 5 min, mounted onto slides using ProLong Diamond Antifade Mountant (Invitrogen), and imaged on a Zeiss LSM 700 confocal microscope with a 63X / 1.4 N.A. objective.

### 4.8. In Vivo Tumor Models

All animal work was approved by the Georgia Tech Institutional Animal Care and Use Committee. Six week old C57BL/6J mice (Jackson Labs) were inoculated with 1×10^5^ B16-OVA cells subcutaneously into the left flank. Tumor burden was monitored until the average tumor volume was approximately 40 mm^3^ prior to adoptive cell transfer. One day before cell infusion, mice were lymphodepleted by irradiation, and OT1 CD8+ T cells were activated as described above for 24 hr. 2×10^5^ activated T cells were transferred via i.v. injection and mice were dosed with 1×10^4^ units recombinant human IL-2 (Peprotech) twice daily for 3 days (6 total injections). Tumor volume was calculated using *V = π/6×L×W×H*. Mice were euthanized when tumor size reached 1.5 cm in any dimension.

## 6. Acknowledgments

This work was funded by an NIH Director’s New Innovator Award (Award DP2HD091793) and by the National Center for Advancing Translational Sciences of the NIH (Award UL1TR000454). S.N.D. is supported by the NSF Graduate Research Fellowships Program (Grant DGE-1650044) and the NSF Integrative Graduate Education and Research Traineeship (Grant DGE-0965945). G.A.K. holds a Career Award at the Scientific Interface from the Burroughs Wellcome Fund. This work was performed in part at the Georgia Tech Institute for Electronics and Nanotechnology, a member of the National Nanotechnology Coordinated Infrastructure, which is supported by the NSF (Grant ECCS-1542174).

## Table of Contents

The accessibility of adoptive T cell therapies is limited by the ability to generate functional T cells. Here we develop synthetic APCs (synAPCs) consisting of liposomes decorated with pMHC molecules for antigen-specific activation of T cells. We show that T cells stimulated by synAPCs exhibit cytotoxic activity and control tumor growth following adoptive transfer *in vivo*.

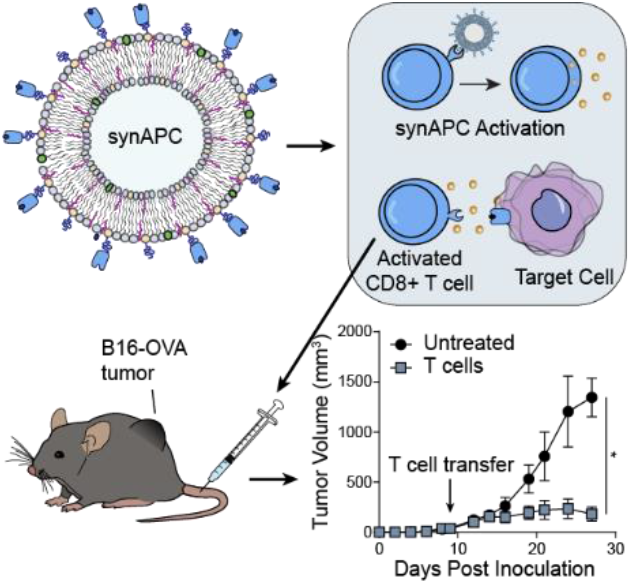

